# BMP2 signaling cooperates with retinoic acid to activate a meiotic-entry transcriptional program in chicken primordial germ cells

**DOI:** 10.64898/2026.07.26.740788

**Authors:** Mohammad Hossein Ghazimoradi, Sofia Corazza, Ali Nemati, Christina Grozou, Cheng Ma, Mohamad Moustafa Ali, Salar Delavari, Ying Xue, Martin Johnsson, Amir Fallahshahroudi, Sara Yousefi Taemeh

## Abstract

The initiation of meiosis in germ cells is largely regulated by extrinsic cues from the gonadal environment, but the logic of these signals remains poorly understood in non-mammalian vertebrates. Retinoic acid has long been considered a principal meiosis-inducing signal, yet recent genetic and reconstitution studies indicate that retinoic acid alone is insufficient. In mice, bone morphogenetic protein 2 cooperates with retinoic acid to establish the oogenic program through the bone morphogenetic protein-responsive transcriptional regulator *Zglp1*, but whether this regulatory logic is conserved beyond mammals is unknown. Here, using cultured chicken (*Gallus gallus*) primordial germ cells, we show that bone morphogenetic protein 2 cooperates with retinoic acid to promote a meiotic-entry transcriptional program. Retinoic acid alone induced a limited retinoic acid-responsive state, whereas combined treatment reduced the primordial germ cell program and activated early meiotic genes. The resulting transcriptome matched the premeiotic-to-meiotic-entry transition of the embryonic ovary in a single-cell atlas of chicken germ cells. We also generated a genome-edited primordial germ cell line carrying an *SYCP3* promoter–green fluorescent protein reporter as a platform for dissecting meiotic-entry signals in culture. Comparative genomic analysis revealed *Gallus*-specific pseudogenization of *ZGLP1*, which is intact in closely related galliform species. Retinoic acid and bone morphogenetic protein may therefore act through a different downstream regulator in chicken.

**Article summary:** Eggs and sperm are produced by meiosis, a specialized cell division that starts during embryonic development. Retinoic acid, a signal derived from vitamin A, was long thought to be enough to start meiosis, but on its own it is not. In mice, a second signal, bone morphogenetic protein 2 (BMP2), works alongside retinoic acid to push germ cells toward the egg-producing program. We asked whether birds use the same combination. Giving both signals to chicken primordial germ cells in culture switched on genes for egg development and for the first steps of meiosis; retinoic acid alone did not. Birds and mammals last shared an ancestor more than 300 million years ago, so the pairing of these two signals appears to be an old feature of vertebrate germ cells. The cells stopped short of completing meiosis, which means other signals are still missing.

**Graphical abstract:** 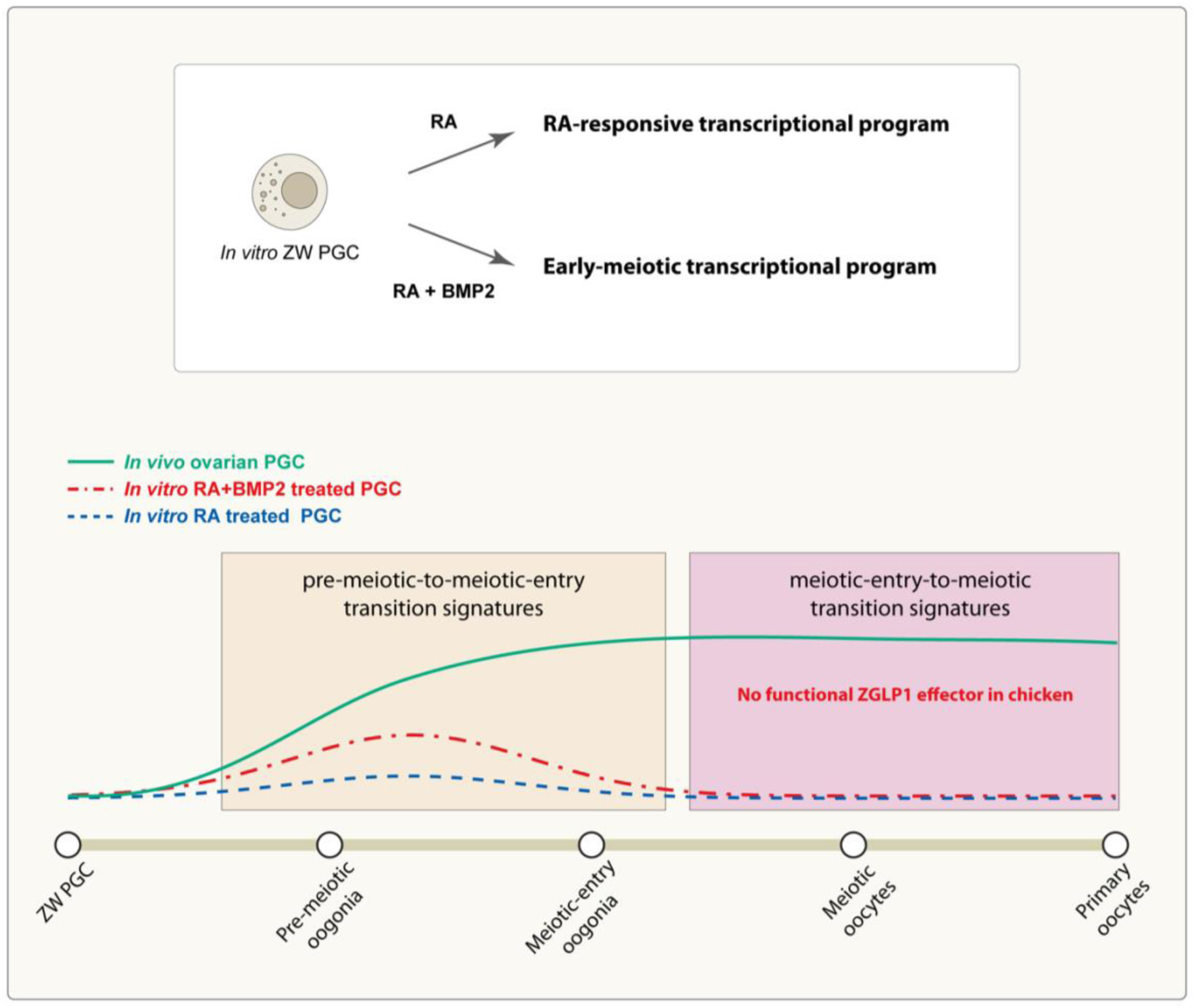

## Introduction

The generation of functional gametes is fundamental to sexual reproduction and requires the timely initiation of meiosis in primordial germ cells (PGCs). In many vertebrates, this developmental transition is tightly regulated by sex-specific extrinsic cues from the gonadal somatic environment. In birds and mammals, female germ cells typically initiate meiosis during embryonic development, whereas male germ cells are prevented from entering meiosis until later developmental stages (Fallahshahroudi et al., 2026; Handel & Schimenti, 2010).

Retinoic acid (RA) has long been considered a principal meiosis-inducing signal in vertebrates. In the classic model, RA promotes meiotic entry in female germ cells by activating the meiotic regulator STRA8, while RA degradation by *CYP26B1* in the fetal testis prevents premature meiotic initiation in males (Anderson et al., 2008; Bowles & Koopman, 2007; Koubova et al., 2006). However, genetic studies have challenged this canonical model. Loss of all nuclear RA receptors or of ALDH1A-dependent RA synthesis does not prevent meiotic initiation in the fetal mouse ovary, although it alters the timing and/or level of *STRA8* expression, implicating additional regulatory pathways (Chassot et al., 2020; Vernet et al., 2020).

Bone morphogenetic protein (BMP) signaling has emerged as one candidate pathway supporting meiotic entry. Studies using mouse *in vitro* germ-cell reconstitution systems have shown that BMP and RA synergistically promote female germ-cell fate, and that RA and its effector STRA8 alone are insufficient to establish a robust oogenic and meiotic program (Miyauchi et al., 2017; Nagaoka et al., 2020) Mechanistically, the BMP-responsive transcription factor zinc finger GATA-like protein 1 (ZGLP1) acts downstream of BMP rather than RA and is required for activation of the oogenic program and meiotic entry (Nagaoka et al., 2020). In line with this model, *in vivo* studies show that BMP signaling is dispensable for the initial induction of *STRA8* but required for correct meiotic progression in fetal mouse germ cells, suggesting that BMP provides an additional input that supports progression beyond STRA8-dependent meiotic initiation (Cheung et al., 2025).

Most mechanistic studies have focused on mammals, and whether BMP provides a complementary input to RA-responsive pathways in non-mammalian vertebrates remains unknown. The chicken (*Gallus gallus*) provides a particularly informative comparative model for addressing this question. RA signaling has been implicated in chicken meiotic onset, with RALDH2-dependent RA synthesis and *CYP26B1*-mediated RA degradation proposed to regulate local RA availability within the developing gonad (Smith et al., 2008; Yousefi Taemeh et al., 2019). Whether a *ZGLP1*-equivalent effector operates downstream of BMP in birds, however, has not been examined.

Chicken PGCs isolated from embryonic blood during migration to the gonad can be propagated extensively *in vitro* while retaining their germ-cell identity and developmental potential. Following transplantation into surrogate host embryos, cultured PGCs can colonize the gonads and ultimately give rise to functional gametes and offspring (Ballantyne et al., 2021; Macdonald et al., 2010). A recent study using this culture system found that exogenous RA induced meiotic-like chromosome morphology and strongly upregulated the RA-degrading enzyme *CYP26A1*; however, canonical meiotic regulators were not significantly induced (Z. Wang et al., 2025). These findings suggest that RA elicits an RA-responsive and potentially self-limiting transcriptional program in chicken PGCs that is insufficient on its own to activate a robust meiotic program.

These observations raise the possibility that additional signals are required to drive meiotic differentiation in chicken PGCs. Given that BMP signaling cooperates with RA/*STRA8*-associated pathways in mouse germ-cell reconstitution systems and supports the correct mitosis-to-meiosis transition in the fetal mouse ovary (Cheung et al., 2025; Miyauchi et al., 2017; Nagaoka et al., 2020), we hypothesized that BMP signaling provides this complementary input in birds.

Here, we used cultured chicken PGCs to test whether BMP2 cooperates with RA to activate a meiotic transcriptional program, benchmarking the induced response against *in vivo* ovarian development and a single-cell atlas of chicken germ-cell differentiation. We also engeneered a *SYCP3*-promoter green fluorescent protein reporter (GFP) PGC line as a platform for dissecting meiotic-entry signals in culture. Finally, we examined the evolutionary conservation of *ZGLP1*, the BMP-responsive effector required for the oogenic program in mammals.

## Results

### Combined RA and BMP2 treatment induces an ovary-like early meiotic transcriptional program in chicken PGCs

To investigate how RA and BMP signaling influence transcriptional programs associated with meiotic differentiation, we performed whole transcriptome analysis of cultured chicken PGCs treated with RA alone or in combination with BMP2. Differential gene expression analysis identified 2,265 genes significantly altered by RA relative to control, including 862 upregulated and 1,403 downregulated genes (FDR-adjusted P < 0.05). Combined RA and BMP2 treatment altered 1,202 genes, of which 307 were upregulated and 895 were downregulated (Table S1 and S2; Fig. 1A). To determine whether these transcriptional responses reflected developmental programs associated with ovarian differentiation, we compared the treatment-induced responses with directionally consistent gene sets derived from a published developmental time series of chicken ovaries from embryonic day 10 to hatching, spanning premeiotic, meiotic-entry, and oocyte-developmental stages (Cardoso-Moreira et al., 2019); Table S1 and S2). Among genes upregulated by each treatment, 29.6% of those induced by RA and BMP2 overlapped with genes upregulated during ovarian development, compared with 20.8% of genes induced by RA alone (Fisher’s exact test, P = 0.0021).

**Figure 1.**
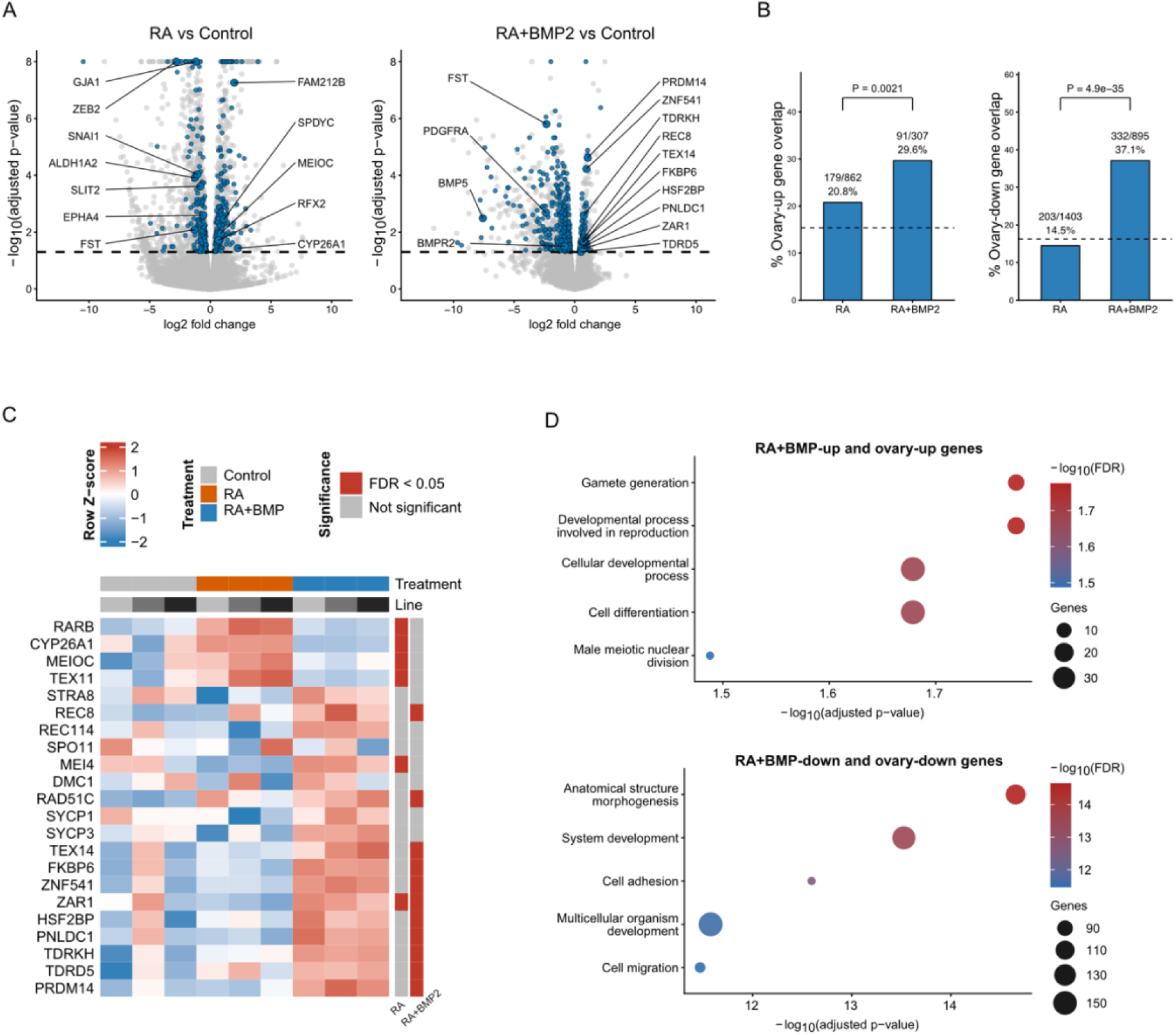
RA and BMP signaling cooperate to induce an ovary-like meiotic differentiation program in chicken PGCs. **A.** Differential gene expression analysis of RA-treated (left) and RA+BMP2-treated (right) PGCs relative to control. Blue points mark treatment-regulated genes (FDR < 0.05) that are directionally matched to ovary developmental genes. Ovary developmental genes were identified by likelihood ratio testing across E10, E14, E17, and P0 ovaries, with direction assigned from E14, E17, and P0 relative to E10 (n=3). **B.** Fraction of RA- or RA+BMP2-regulated genes that are directionally matched to ovary developmental genes as defined in A, divided by the number of upregulated or downregulated genes in each group. Dashed lines indicate the expected overlap based on the background gene set; P values are from Fisher’s exact tests comparing RA and RA+BMP2. **C.** Row-scaled expression of selected RA-response, meiotic, recombination, and germ-cell differentiation genes across control, RA, and RA+BMP2 samples; side bars indicate genes differentially expressed at FDR < 0.05. **D.** Gene Ontology enrichment for directionally matched RA+BMP-upregulated genes (left) and RA+BMP-downregulated genes (right). Point size indicates gene count; color denotes −log10(FDR).

This difference was more pronounced among downregulated genes: 37.1% of genes downregulated by RA and BMP2 overlapped with genes downregulated during ovarian development, compared with 14.5% following RA treatment alone (Fisher’s exact test, P = 4.9 × 10⁻³⁵; Fig. 1B). Thus, combined RA and BMP2 treatment produced a transcriptional response that was more concordant with ovarian developmental regulation, particularly among genes repressed during ovary development.

RA alone predominantly activated canonical RA-responsive genes, including *RARB* and *CYP26A1*, along with a broader differentiation signature comprising *MYO1C*, *HNF1B*, and *SOX5*. This response was concordant with the transcriptional changes reported by previous work (Z. Wang et al., 2025), supporting the consistency of our experimental system with earlier findings. However, induction of meiosis-associated genes was limited, and *STRA8*, *REC8*, *RAD51C*, and *TEX14* were not significantly induced. In contrast, combined RA and BMP2 treatment significantly upregulated *REC8*, *RAD51C*, *TEX14*, *ZAR1*, and *ZNF541*, together with the piRNA-pathway components *TDRKH*, *TDRD5*, *PNLDC1*, and *FKBP6*. Nevertheless, later meiotic-prophase markers, including *DMC1*, *SPO11*, and *SYCP1*, did not meet the significance threshold (FDR < 0.05), whereas *SYCP3* showed only a trend toward induction (Fig. 1C; Table S1 and S2). Combined treatment therefore activated transcriptional components of an oogenic and early meiotic program but did not establish a complete meiotic-prophase state.

Consistent with these gene-level changes, genes upregulated by combined RA and BMP2 treatment were enriched for reproductive and gametogenesis-related processes, including gamete generation and reproductive development. Conversely, downregulated genes were enriched for broader developmental and morphogenetic processes, including anatomical structure morphogenesis, cell adhesion, multicellular-organism development, and cell migration (Fig. 1D). Several genes induced by the combined treatment were also activated by *Zglp1*overexpression in mouse PGC-like cells, in which RA promotes further maturation of the ZGLP1-induced program (Nagaoka et al., 2020).

Together, these results indicate that RA primarily activates an RA-responsive transcriptional program, whereas combined RA and BMP2 treatment shifts chicken PGCs toward a more ovary-like state with oogenic and early meiotic features. However, the limited induction of later meiotic-prophase markers indicates that the combined treatment is insufficient to establish a complete meiotic program.

### RA and BMP2-associated transcriptional signatures peak at the premeiotic-to-meiotic-entry transition

Having shown that combined RA and BMP2 treatment produces a transcriptional response that is more concordant with ovarian development than the response to RA alone, we next examined where these concordant gene-expression signatures were represented along the female germ-cell trajectory *in vivo*. We first defined two gene sets by identifying genes that changed in the same direction in RA+BMP2-treated PGCs and during ovarian development. The induced signature comprised genes upregulated in both datasets, whereas the repressed signature comprised genes downregulated in both datasets. Because bulk ovary transcriptomes contain signals from both germ and somatic cells, we assessed these predefined gene sets in a published single-cell transcriptomic atlas of female chicken germ-cell development spanning embryonic day 2.5 (E2.5) to one week post-hatch (Rengaraj et al., 2022a). This allowed us to determine the germ-cell states in which each bulk-derived signature was most strongly represented.

Female germ cells were distributed along a developmental trajectory in UMAP space, progressing from PGCs through pre-meiotic oogonia, meiotic-entry oocytes, and meiotic oocytes to primary oocytes (Fig. 2A). These annotations were supported by established stage-associated expression patterns. PGCs expressed high levels of *NANOG*, pre-meiotic oogonia were enriched for *REC8*, and meiotic-entry oocytes showed strong *STRA8* expression. Meiotic oocytes expressed recombination-associated genes, including *SPO11* and *DMC1*, whereas primary oocytes were characterized by high expression of the oocyte-development factor *LHX8* (Fig. 2B).

**Figure 2.**
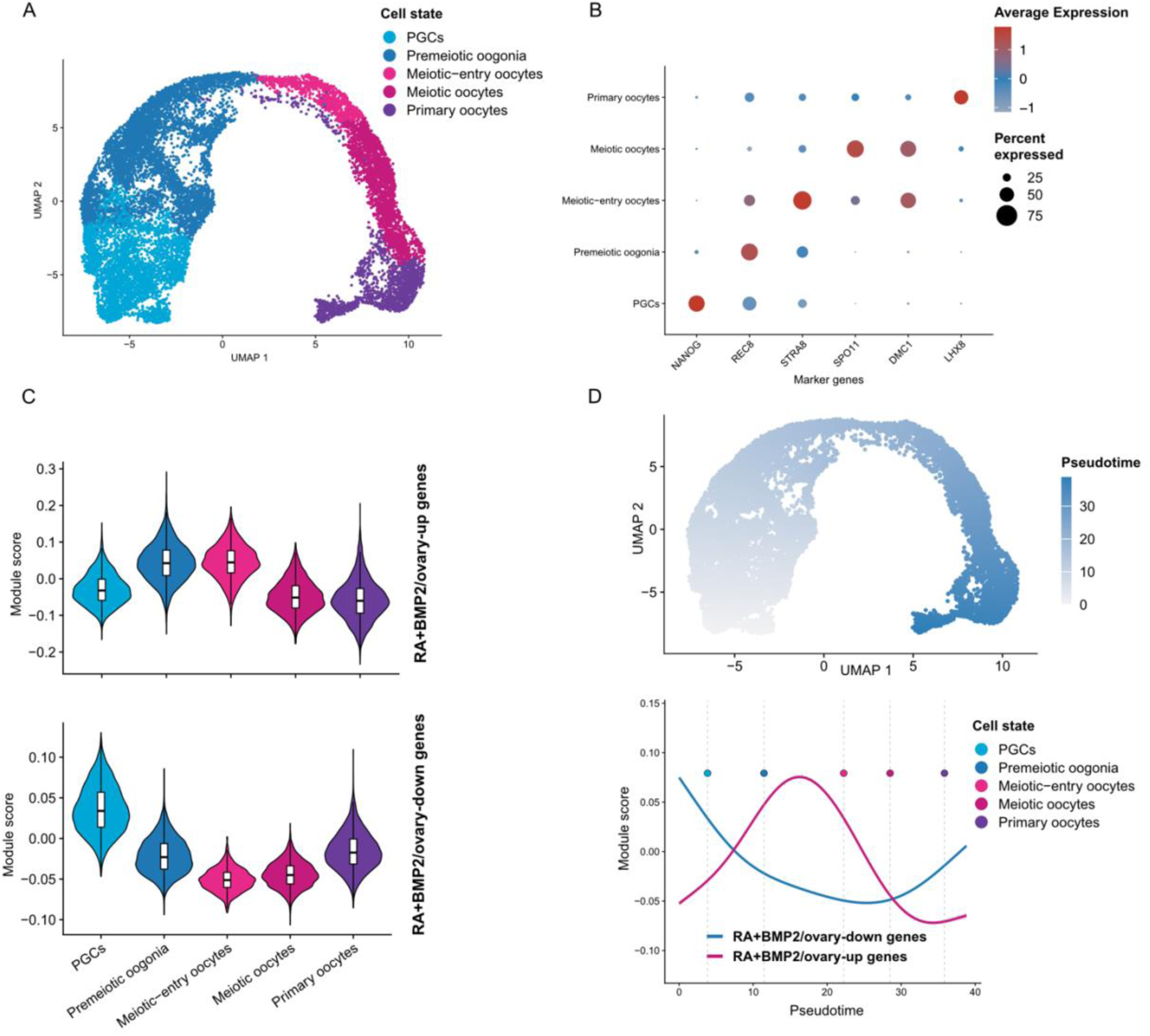
RA and BMP2-associated gene programs peak at the pre-meiotic-to-meiotic-entry transition. A. UMAP of female chicken germ cells colored by annotated germ-cell state. **B**. Expression of representative germ-cell state markers across annotated cell states. Dot size indicates the percentage of cells expressing each gene, and color indicates average scaled expression. **C.** Module scores for directionally matched RA+BMP2 gene sets across germ-cell states. The upper violin plot shows RA+BMP2-upregulated genes that also increase during ovarian development, and the lower violin plot shows RA+BMP-downregulated genes that also decrease during ovarian development. Boxplots indicate the median and interquartile range. **D.** Pseudotime projected onto the UMAP embedding (top) and smoothed module-score dynamics for the same directionally matched RA+BMP2 gene sets along pseudotime (bottom). Dashed lines and colored points indicate the median pseudotime of each annotated germ-cell state.

We used Seurat module scoring to quantify the expression of the concordantly induced and repressed gene sets across this trajectory. Module scores represent the average expression of each gene set relative to that of expression-matched control genes. The induced signature showed its highest scores in pre-meiotic oogonia and meiotic-entry oocytes. In contrast, the repressed signature showed its highest scores in PGCs, followed by a progressive decline across the pre-meiotic and meiotic-entry states (Fig. 2C). Pseudotime analysis revealed similar reciprocal dynamics: the repressed signature was highest at the beginning of the trajectory, whereas the induced signature increased after the PGC state, peaked near the transition from pre-meiotic oogonia to meiotic-entry oocytes, and declined in later oocyte states (Fig. 2D). These patterns are consistent with combined RA and BMP2 treatment suppressing an early PGC-like transcriptional program while activating genes normally associated with the transition from the pre-meiotic to the meiotic-entry state.

### A *SYCP3* promoter-EGFP reporter monitors partial meiotic activation in cultured chicken PGCs

Although RA and BMP2 treatment promoted a meiotic-entry transcriptional program, several later meiotic prophase markers were not fully induced under standard culture conditions, suggesting that cultured PGCs may require further optimization of culture conditions to progress beyond the pre-meiotic-to-meiotic-entry state. Because these conditions are formulated to sustain long-term proliferation and germ-cell identity, we reasoned that self-renewal signals in the medium might oppose meiotic differentiation. To directly monitor progression toward meiotic prophase, we generated a *SYCP3*pro-EGFP reporter ZW PGC line by identifying and targeting a genetic safe-harbor locus (GSH) (Table S3 and Fig. 3A). Because complete withdrawal of FGF and Activin was not tolerated—PGCs died within 72 h—we used this line to test whether reducing, rather than removing, growth-factor support while increasing RA and BMP2 signaling could enhance meiotic induction.

**Figure 3.**
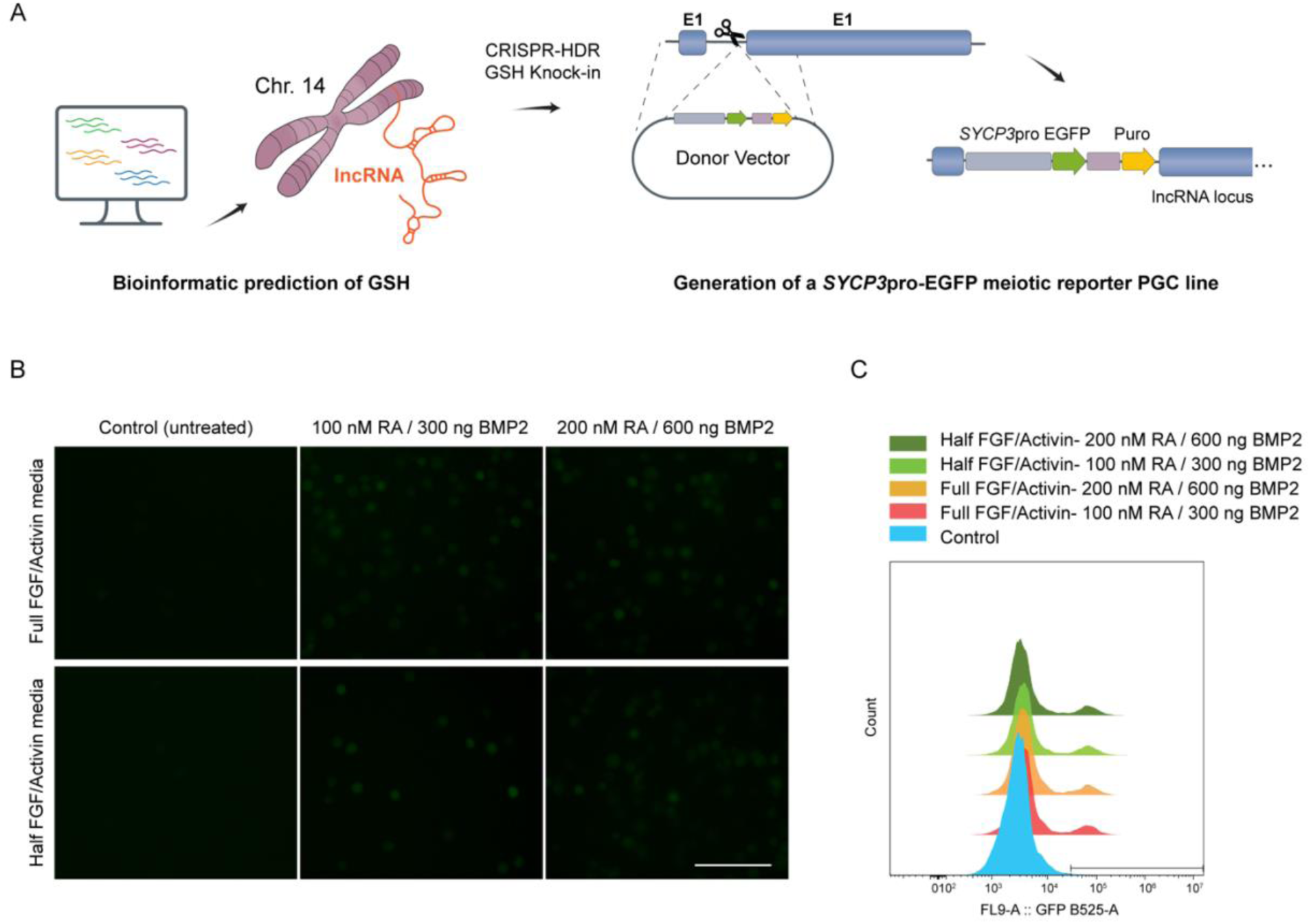
Generation and use of a *SYCP3*pro-EGFP ZW PGC reporter line to monitor meiotic activation. **A.** Schematic overview of the strategy used to generate the *SYCP3*pro-EGFP meiotic reporter PGC line. A candidate genetic safe-harbor locus was identified by bioinformatic screening of highly expressed and stably expressed long non-coding RNA loci. The *SYCP3*pro-EGFP reporter cassette, together with a puromycin-resistance selection marker, was inserted into an intronic region of the selected chromosome 14 lncRNA locus by CRISPR/Cas9-mediated homology-directed repair (HDR) knock-in. **B.** Fluorescence microscopy of *SYCP3*pro-GFP reporter PGCs after RA and BMP2 treatment under full or reduced growth-factor conditions. Cells were cultured in full FGF/Activin medium or half FGF/Activin medium and treated with either 100 nM RA and 300 ng/mL BMP2 or 200 nM RA and 600 ng/mL BMP2. Untreated cells served as controls. Scale bar, 100 µm. **C.** Flow cytometry analysis of GFP fluorescence in *SYCP3*pro-EGFP reporter PGCs under the same treatment conditions.

RA and BMP2 treatment increased GFP fluorescence in the SYCP3pro-EGFP reporter line compared with untreated control cells, consistent with activation of the reporter during meiotic induction. However, the GFP signal remained modest, in agreement with the limited increase in SYCP3 expression observed in our RNA-sequencing data (log2 fold change = 0.8, FDR-adjusted P < 0.1). Reducing FGF/Activin support, either alone or together with increased RA and BMP2 concentrations, did not further enhance reporter fluorescence. Consistently, flow cytometry analysis under the same treatment conditions showed increased GFP fluorescence in RA+BMP2-treated cells relative to untreated controls, but no clear difference between full and half FGF/Activin medium or between the tested RA/BMP2 doses (Fig. 3B,C). These results suggest that additional ovarian niche-derived cues, beyond RA and BMP2, may be required to support more complete meiotic differentiation in cultured chicken PGCs.

### Ovarian somatic and germline cells form a complex intercellular signaling network

To identify candidate niche-derived signals that might support germ-cell differentiation beyond the partial meiotic-entry state induced by RA and BMP2, we analyzed a published E17–E19 chicken ovary scRNA-seq dataset containing both germline and non-germline populations (Trost *et al*., 2025; Fig. 4A). This developmental window encompasses meiotic progression *in vivo*, during which female germ cells transition from pre-meiotic stages into early meiotic prophase I. Using CellChat (Jin *et al*., 2024), we inferred ligand– receptor communication among these populations and identified 3,373 significant ligand–receptor interactions across cell-type pairs, revealing a dense intercellular communication network (Fig. 4B). The global interaction map showed extensive communication among PGCs, pre-meiotic oogonia, meiotic oogonia, and surrounding non-germline populations, including pre-granulosa, theca, mesenchymal, endothelial, and coelomic epithelial cells (Fig. 4A, B).

**Figure 4.**
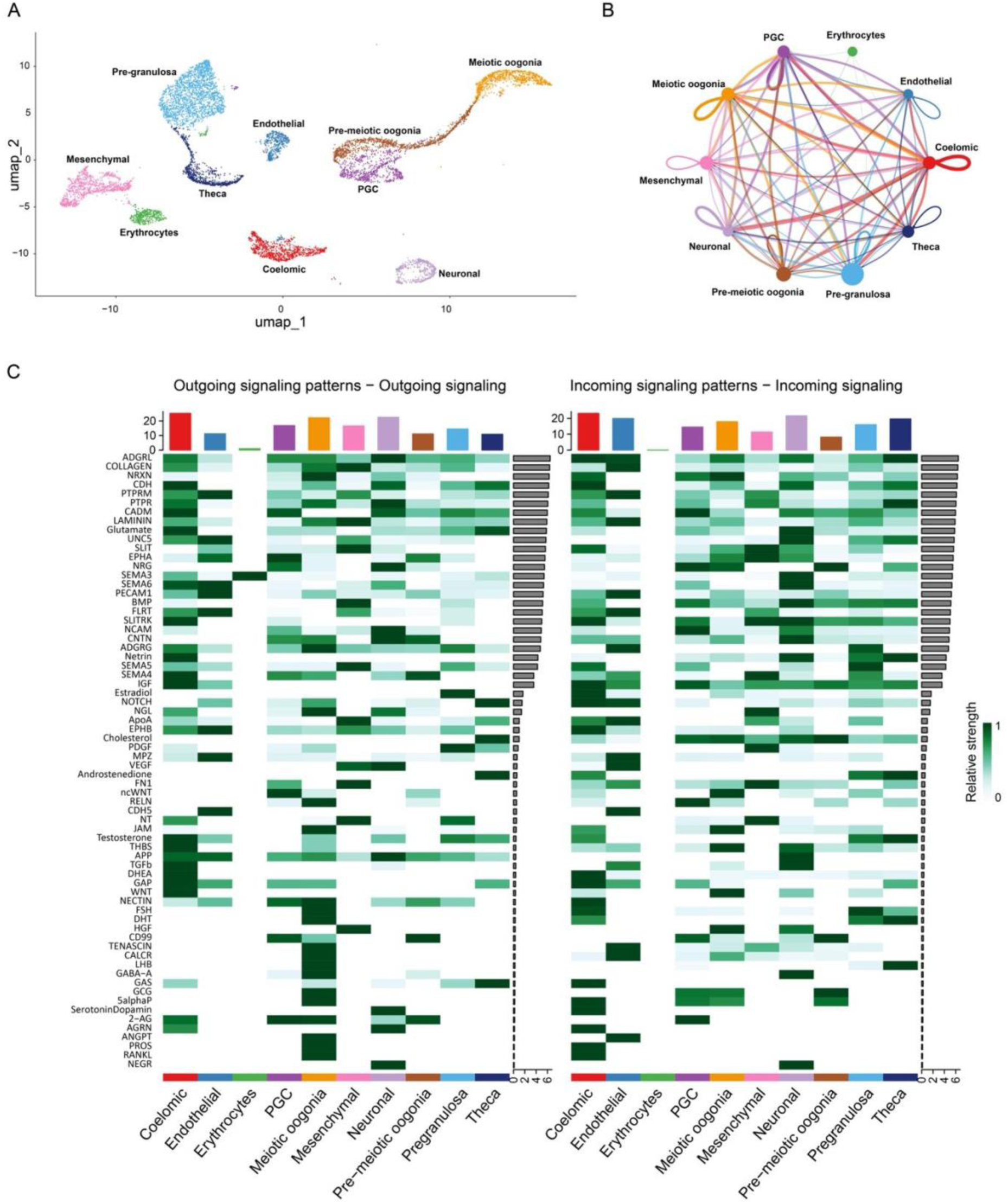
Intercellular signaling networks across ovarian somatic and germline compartments in the E17–E19 chicken ovary. **A.** UMAP projection of annotated cell populations from the E17–E19 chicken ovary scRNA-seq dataset (n = 8,586 cells), colored by cell type. Non-germline populations comprise coelomic epithelial, pre-granulosa, theca, mesenchymal, endothelial, neuronal, and erythrocyte cells, whereas germline populations comprise PGCs, pre-meiotic oogonia, and meiotic oogonia. **B.** CellChat-inferred global communication network among ovarian cell populations. The network displays a combined interaction score calculated as the mean of the scaled interaction count and scaled interaction strength. Node size is proportional to the number of cells in each population, edge width represents the combined interaction score, and edge color indicates the sending cell population. Self-loops indicate inferred autocrine signaling. **C.** Heatmaps of outgoing and incoming signaling patterns across germline and non-germline populations. Rows represent inferred signaling pathways, and columns represent cell populations. Color intensity indicates relative signaling strength, scaled from 0 to 1. Bars above the heatmaps summarize the total outgoing or incoming signaling strength of each cell population, whereas grey bars to the right summarize the total signaling strength of each pathway.

Analysis of outgoing and incoming signaling patterns showed that germline populations acted as both signal senders and receivers (Fig. 4C). Among the germline populations, meiotic oogonia and PGCs exhibited the strongest overall outgoing signaling activity, whereas pre-meiotic oogonia showed a comparatively weaker profile. Non-germline populations, including pre-granulosa, theca, mesenchymal, coelomic epithelial, and endothelial cells, also displayed distinct incoming and outgoing signaling patterns, consistent with a complex ovarian signaling niche during meiosis. The inferred communication pathways included BMP, TGF-β, canonical and non-canonical WNT, NRG, PDGF, NOTCH, EPHA/EPHB, SLIT, SEMA, SLITRK, PTPR, NCAM, CNTN, LAMININ, COLLAGEN, TENASCIN, and other adhesion- and extracellular-matrix-associated pathways (Fig. 4C). Several of these pathways showed incoming activity in one or more germline populations, suggesting that germ cells receive multiple extrinsic inputs beyond RA and BMP2. The inferred incoming and outgoing signaling profiles also differed among PGCs, pre-meiotic oogonia, and meiotic oogonia, indicating that communication with the ovarian niche changes during germ-cell differentiation (Fig. 4C and Table S4). Together, these results support a model in which meiotic differentiation in chicken germ cells is shaped by a broader ovarian signaling environment and identify BMP/TGF-β, WNT, NRG/PDGF, and extracellular-matrix-associated pathways as candidates for improving in vitro meiotic induction.

### *ZGLP1* is retained as a pseudogenized relict in the chicken genome but remains intact in galliform sister species

ZGLP1 functions downstream of BMP signaling and promotes oogenic differentiation and meiotic entry in the mouse fetal ovary. We therefore investigated whether *ZGLP1* is conserved in chicken using protein homology searches, genome-wide nucleotide searches, and synteny analysis across vertebrate assemblies (Methods; Table S5).

We examined the Huxu telomere-to-telomere and GRCg6a chicken assemblies. In both assemblies, *ZGLP1*-derived sequence was identified on chromosome 30 at the conserved syntenic position observed in other birds (Fig. 5A). The chicken locus retained the N-terminal portion of the canonical GATA-type zinc-finger domain with complete sequence identity to the corresponding region of *Lagopus muta ZGLP1*. However, premature stop codons disrupted the remainder of the zinc-finger domain, and the locus contained a pronounced proline-rich expansion, indicating that chicken *ZGLP1* is pseudogenized rather than protein coding.

**Figure 5.**
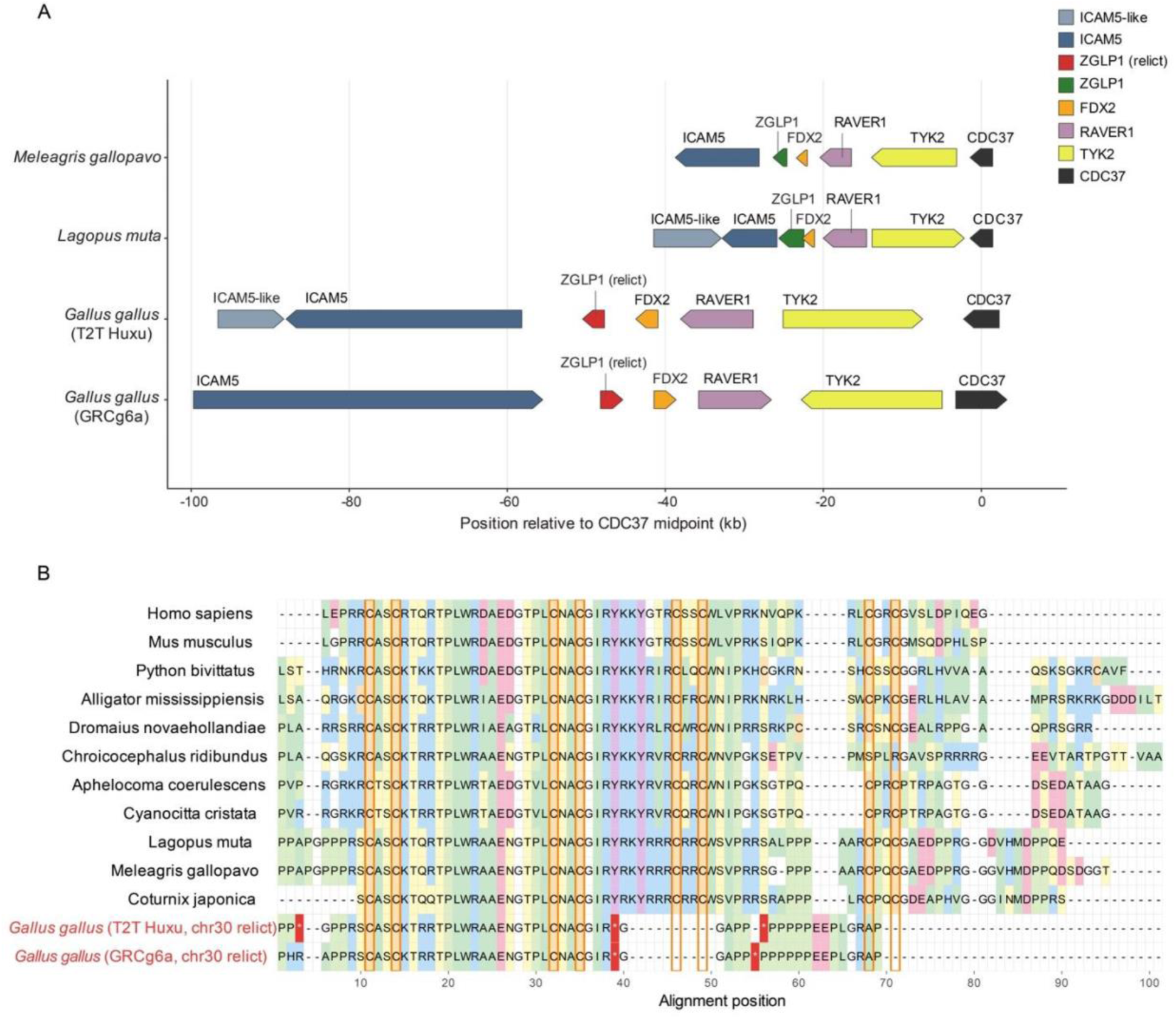
*ZGLP1* persists as a pseudogenized relict on chicken chromosome 30. **A.** Conserved synteny of the *ICAM5*–*ZGLP1*–*FDX2*– RAVER1–TYK2–CDC37 interval in two galliform species with intact *ZGLP1, Meleagris gallopavo* (NW_028147023.1) and *Lagopus muta* (NW_026040201.1), and two chicken genome assemblies containing a ZGLP1 relict: *Gallus gallus* T2T Huxu (CP100584.2:149,802–152,582; tblastn E = 1.70 × 10⁻⁴³) and GRCg6a (NC_028739.2:219,654–222,480; E = 4.89 × 10⁻⁴⁰). Searches were performed using the L. muta ZGLP1 protein XP_048788870.1 as the query. Arrows indicate transcriptional orientation, and genomic positions are shown in kilobases relative to the midpoint of *CDC37.* Intact *ZGLP1* genes are shown in green and chicken relicts in red. **B.** MAFFT L-INS-i alignment of the ZGLP1 zinc-finger region from 13 sequences representing 12 vertebrate species. Galliform sequences include *L. muta*, *M. gallopavo*—in which ZGLP1 is annotated as part of the LOC104913635 fusion protein—*Coturnix japonica*, reconstructed from GCA_001577835.2, and the relict sequences from the two chicken assemblies, shown in red. Zinc-coordinating cysteines are outlined in orange, and in-frame stop codons are indicated by red asterisks. The chicken relicts retain the N-terminal CASCKTRRTPLWRAAENGTPLCNACGIR motif but are truncated downstream and contain a proline-rich expansion, whereas the three other galliform species retain intact zinc-finger regions.

To determine whether this loss occurred more broadly within Galliformes, we examined ZGLP1 sequences from closely related species. Intact zinc-finger sequences without premature stop codons were recovered from the examined galliform species, including turkey, in which ZGLP1 was incorporated into an incorrectly annotated fusion protein. Alignment across representative vertebrates showed conservation of the eight zinc-coordinating cysteine residues in these species, whereas both chicken assemblies retained only the N-terminal half of the domain (Fig. 5B; Table S5; File S1).

Together, the conserved syntenic position and sequence identity establish the chromosome 30 locus as the chicken *ZGLP1* relict and indicate that *ZGLP1* pseudogenization occurred specifically in the Gallus lineage rather than throughout Galliformes.

## Discussion

Here, we show that adding BMP2 to RA shifts cultured chicken PGCs toward an ovary-like oogenic and early meiotic transcriptional state. This combined RA and BMP2 response showed stronger concordance with ovarian developmental gene-expression changes than RA alone and was most strongly represented *in vivo* at the pre-meiotic-to-meiotic-entry transition. However, several later meiotic-prophase markers were not fully induced, indicating that RA and BMP2 promotes only partial meiotic activation. Moreover, adjusting growth-factor concentrations and increasing RA/BMP2 doses did not further advance meiotic progression in our meiotic reporter PGC line. This incomplete response is consistent with our ligand-receptor analysis, which revealed a complex ovarian signaling niche and suggests that additional extrinsic cues may be required beyond RA and BMP2. Together, these findings suggest that RA- and BMP-responsive pathways may interact to promote early meiotic differentiation in chicken PGCs, as described in mouse germ-cell reconstitution systems. Because our comparative genomic analysis places *ZGLP1* as a pseudogenized relict, chickens likely use a divergent or alternative BMP-responsive regulator to link BMP signaling to activation of the oogenic and meiotic-entry program.

Our RA-alone results are consistent with the recent RNA-seq analysis of RA-treated chicken PGCs by (Z. Wang et al., 2025), which showed strong induction of *CYP26A1* and other differentiation-associated genes, including *MYO1C*, *HNF1B*, and *SOX5*. Similarly, we found that RA alone predominantly activated canonical RA-responsive and differentiation-associated genes but had limited effects on meiosis-associated genes. Importantly, Wang, Z. et al also reported limited induction of *STRA8*, supporting the idea that RA alone does not establish a robust meiotic transcriptional program in cultured chicken PGCs (Fig. 1A, B).

This interpretation is also consistent with our previous *ex vivo* analysis of embryonic chicken gonads, which showed that meiotic initiation is not tightly coupled to *STRA8* transcript levels and requires additional gonad-associated or mesonephros-derived cues (Yousefi Taemeh et al., 2019). Our current study extends these findings by showing that the addition of BMP2 broadens the RA response and significantly induces early meiotic and oogenic genes, including *REC8*, *RAD51C*, *TEX14*, *ZAR1*, and *ZNF541* (Fig. 1C). This transcriptomic shift was further supported by gene ontology enrichment analysis, which showed that the combined treatment activates genes associated with gamete generation and reproductive differentiation while repressing genes linked to cell migration, cell adhesion, anatomical structure morphogenesis, and broader developmental programs (Fig. 1D). This cooperative effect suggests that, in chicken PGCs, RA responsiveness alone is insufficient for meiotic transcriptional priming. Rather, BMP2 provides an additional developmental signal that supports the acquisition of an early oogenic-like state, but not the establishment of a fully developed oogenic program. This is consistent with mouse models showing that RA–*Stra8* signaling is important for meiotic initiation (Anderson et al., 2008; Bowles & Koopman, 2007) but is insufficient to establish the full oogenic and meiotic program. BMP signaling, partly through *Zglp1*, provides an additional input required for female germ-cell fate specification and proper meiotic progression (Cheung et al., 2025; Miyauchi et al., 2017; Nagaoka et al., 2020).

In mice, ZGLP1 acts downstream of BMP signaling rather than RA and is required to activate the oogenic program and support meiotic entry (Nagaoka et al., 2020). Related roles of ZGLP1/*Zglp1* have been reported in other vertebrates, including zebrafish, where *zglp1* deficiency disrupts ovarian development and causes female-to-male sex reversal (Y. Wang et al., 2025). In zebra finch, the only identifiable *ZGLP1* copy is confined to the germline-restricted chromosome after duplication from chromosome 30 and loss from the standard A chromosomes (Ruiz-Ruano et al., 2025). Our comparative-genomic analysis extends these observations by placing a *ZGLP1* pseudogenization event on the Gallus lineage (Fig. 5). Because two independent chicken assemblies recover the same relict with negligible ambiguous-base content, the pseudogenized state cannot be attributed to assembly gaps. Our data therefore indicate that chickens have lost a functional *ZGLP1* and likely engage a distinct or lineage-specific BMP-responsive regulator to couple BMP signalling to the oogenic and meiotic-entry program that we observe active in cultured chicken PGCs; identifying and functionally testing that regulator is an important direction for future work.

We also observed upregulation of several piRNA pathway-associated genes following combined RA and BMP2 treatment, including *TDRKH*, *TDRD5*, *PNLDC1*, and *FKBP6* (Fig. 1C). The piRNA pathway is a conserved, germline-enriched small RNA silencing mechanism that safeguards genome integrity by suppressing transposable elements (Czech et al., 2018; Tóth et al., 2016). In chicken, core PIWI-like proteins (such as CIWI and CILI) are crucial for suppressing chicken-specific transposable elements, particularly chicken repeat 1 (CR1) LINE elements, during critical windows of germline development (Chang et al., 2018; Kim et al., 2012) and for the protection of PGCs from transposon-associated DNA damage (Rengaraj et al., 2014). Therefore, the induction of piRNA pathway components observed here may indicate that combined RA and BMP2 treatment initiates a broader germline maturation program in chicken PGCs, consistent with a developmental shift toward an early oogenic-like state. However, because our current transcriptomic analysis measured changes in expression of piRNA pathway-associated genes rather than mature piRNAs themselves, functional activation of this pathway will require future small RNA profiling together with direct assessment of transposable element expression in treated cells.

Despite the induction of early meiotic and oogenic genes, markers of meiotic-prophase progression were not significantly induced, suggesting that RA and BMP2 alone are insufficient to drive full meiotic entry in cultured chicken PGCs. Reducing FGF and Activin did not further enhance SYCP3pro-EGFP reporter activity, indicating that partial attenuation of self-renewal signaling is also insufficient to promote progression. This may reflect residual FGF/Activin activity or a requirement for additional signals from the ovarian niche. Our ligand–receptor analysis supports the latter possibility. In vivo, E17–E19 chicken ovaries contain a complex signaling network in which germ cells interact with multiple somatic populations and receive inputs from BMP/TGF-β, non-canonical WNT, NRG, PDGF, Notch, adhesion, and extracellular-matrix pathways (Fig. 4A–C). RA and BMP2 therefore likely represent only part of the signaling environment required for meiotic differentiation. Because the inferred signals change as germ cells progress from PGCs toward meiosis, successful differentiation may require sequential, stage-specific cues rather than a fixed culture condition. The enrichment of adhesion and ECM pathways further suggests that somatic-cell contact and matrix support, which are absent from suspension culture, may facilitate meiotic progression.

In addition to these extrinsic requirements, developmentally regulated chromatin states or other epigenetic constraints may limit activation of downstream meiotic-prophase genes in cultured PGCs (Miyauchi et al., 2017; Nagaoka et al., 2020). This interpretation is consistent with the relatively late onset of germ-cell differentiation in the chicken ovary, where meiotic germ cells are first detected around embryonic day 15.5 (Smith et al., 2008), and with evidence that gonad-associated or mesonephros-derived cues contribute to meiotic initiation (Yousefi Taemeh et al., 2019).

The GSH-targeted SYCP3pro-EGFP reporter line provides a tractable platform for testing these paracrine and structural cues, including WNT pathways, alternative BMP ligands, somatic-cell-conditioned media, and epigenetic modulators, through combinatorial in vitro screening and in vivo germline transplantation. Furthermore, the identified lncRNA locus expands the repertoire of available chicken GSH sites beyond the intergenic loci described previously (Dehdilani et al., 2023).

Overall, our findings support a model in which RA establishes an RA-responsive differentiation state in chicken PGCs, whereas BMP2 cooperates with RA to shift the transcriptional trajectory toward an early oogenic and meiotic program. However, the limited induction of meiotic-prophase genes indicates that RA and BMP2 are sufficient for meiotic priming but not for complete meiotic progression. Efficient *in vitro* meiosis in avian germ cells may therefore require the coordinated delivery of stage-specific signaling cues, appropriate chromatin competence, and structural support from the ovarian niche. Defining these additional regulatory and microenvironmental requirements will be an important next step toward establishing robust *in vitro* gametogenesis systems in chicken and extending such approaches beyond mammalian models.

## Materials and methods

### Chicken PGC derivation, sexing, and RA/BMP2 treatment

To establish PGC lines, blood was collected from Lohmann Brown chicken embryos at Hamburger– Hamilton stage Stages 14 to 16 (E2.5). Ten independent PGC lines were derived and cultured according to previously described methods (Dehdilani, Yousefi Taemeh, et al., 2023; Yousefi Taemeh et al., 2025). The sex of each line was determined by PCR using two primer sets: one targeting a W-chromosome-specific gene and one targeting an autosomal control gene, as described previously (Fallahshahroudi et al., 2025) Three female PGC lines were selected for experimental treatments.

For meiotic induction experiments, cultures were treated for 7 days with either retinoic acid (RA) alone at a final concentration of 100 nM, or with a combination of 100 nM RA and 300 ng/mL bone morphogenetic protein 2 (BMP2). These concentrations were based on conditions previously used for meiotic induction in mouse germ-cell reconstitution systems (Miyauchi et al., 2017). Medium containing the respective factors was fully replaced every 3 days. Untreated cells maintained under standard culture conditions served as controls.

### RNA extraction, library preparation, and sequencing

Total RNA was extracted from approximately 1 × 10⁷ cultured PGCs per sample using TRIzol Reagent (Thermo Fisher Scientific) according to the manufacturer’s instructions. RNA integrity was assessed using an Agilent 2100 Bioanalyzer, and all samples had an RNA Integrity Number (RIN) > 8 before library preparation.

Libraries were prepared from three biological replicates per treatment group using the NEBNext Ultra II Directional RNA Library Prep Kit for Illumina (New England Biolabs), following poly(A) mRNA enrichment. Polyadenylated RNA was isolated using oligo(dT) magnetic beads, thermally fragmented, and reverse-transcribed to generate strand-specific cDNA libraries. Libraries were then subjected to end repair, adapter ligation. Library preparation and sequencing were performed by Novogene Co., Ltd. Libraries were sequenced on an Illumina NovaSeq X Plus platform using paired-end 150 bp chemistry, generating approximately 30 million reads per sample.

For the control condition, two technical replicates were generated for each biological line by independently expanding the same line under untreated conditions, followed by separate RNA extraction, library preparation, and sequencing.

### RNA-seq data processing and differential expression analysis

The chicken reference genome bGalGal1.mat.broiler.GRCg7b and corresponding GTF annotation file were obtained from Ensembl release 115. Raw reads from each library were aligned to the reference genome using STAR v2.7.11b (Dobin et al., 2013). STAR genome indices were generated using the reference genome and GTF annotation, and reads were aligned in an annotation-aware manner. Gene-level read counts were generated using the STAR quantMode GeneCounts option. Technical replicates were averaged for each line before downstream analysis.

Gene count matrices, together with sample metadata and gene annotation information, were used to construct a RangedSummarizedExperiment object in DESeq2 v1.52.0 (Love et al., 2014) for downstream analysis. Protein-coding genes were retained based on the GTF annotation before differential expression analysis. Size factors were estimated using DESeq2, and differential expression was tested using a design formula that included PGC line as a blocking factor: ∼ line + treatment. This design accounts for paired biological variation among the three independent PGC lines. Differential expression was assessed for RA versus control and RA and BMP versus control using DESeq2 contrast testing. Genes were considered differentially expressed at Benjamini-Hochberg false discovery rate (FDR) corrected P-values < 0.05.

### Gene ontology enrichment analysis

GO Biological Process enrichment analysis was performed using differentially expressed genes from the RA versus control and RA and BMP versus control comparisons. Chicken genes were mapped to human one-to-one orthologs using BioMart, and only genes with valid human Ensembl IDs and gene symbols were retained. For each comparison, the enrichment background was defined as all detected genes with mapped human orthologs.

Differentially expressed genes were selected at adjusted *P* < 0.05 and separated into upregulated and downregulated gene sets based on the sign of the DESeq2 log_2_ fold change. GO enrichment was performed using gprofiler2::gost() with human gene symbols and organism = "hsapiens", using the corresponding custom background for each comparison. Terms were considered enriched at FDR < 0.05.

For visualization, enriched GO Biological Process terms were ranked by adjusted enrichment *P* value, and selected top terms were displayed as dot plots. The x-axis represents −log_10_ adjusted enrichment *P* value, and point size indicates the number of genes overlapping each GO term. Broad or uninformative root terms were excluded from visualization.

### Integration of bulk ovarian RNA-seq with the single-cell germ-cell atlas

Public chicken ovary RNA-seq data from E10, E14, E17, and P0, with two biological replicates per stage, were obtained from Cardoso-Moreira et al. (2019). FASTQ files were generated from Sequence Read Archive accessions using fasterq-dump from the SRA Toolkit v3.4.1. Reads were processed using the same reference genome and annotation as the experimental RNA-seq data, aligned with STAR, and quantified at the gene level using --quantMode GeneCounts.

Differential expression across ovarian development was assessed using a DESeq2 likelihood-ratio test. Developmental stage was modeled as a categorical factor, with E10 as the reference level. The full model, ∼ stage, was compared with the reduced intercept-only model, ∼ 1. Protein-coding genes with an LRT-adjusted P < 0.05 were considered developmentally regulated.

To assign directionality, pairwise log2 fold changes were extracted for E14 versus E10, E17 versus E10, and P0 versus E10. Genes were classified as ovary up-only when they were significant in the likelihood-ratio test and showed a positive log2 fold change in at least one post-E10 comparison without showing a negative log2 fold change in any comparison. Genes were classified as ovary down-only using the opposite criterion. Genes showing both positive and negative log2 fold changes across post-E10 comparisons were classified as mixed and excluded from direction-matched analyses.

Direction-matched overlap was assessed by comparing treatment-upregulated genes with ovary up-only genes and treatment-downregulated genes with ovary down-only genes. The background gene set was defined as the intersection of genes included in the RA, RA+BMP2, and ovarian developmental analyses. Differences in overlap between the RA and RA+BMP2 conditions were tested using Fisher’s exact test.

To connect these bulk RNA-seq analyses with the single-cell atlas, we defined directionally concordant gene signatures from the RA+BMP2 treatment and ovarian developmental datasets. The induced signature comprised genes that were significantly upregulated by RA+BMP2 treatment, significant in the ovarian developmental likelihood-ratio test, and classified as ovary up-only. The repressed signature comprised genes that were significantly downregulated by RA+BMP2 treatment, significant in the ovarian developmental likelihood-ratio test, and classified as ovary down-only. Mixed-direction genes were excluded.

Public single-cell RNA-seq data spanning chicken germ-cell development were obtained from Rengaraj et al. (2022) and reprocessed from raw reads. Reads were mapped to the bGalGal1.mat.broiler.GRCg7b chicken reference genome using STARsolo, and gene-count matrices were generated for downstream analysis. Cells were processed in Seurat v5.5.0 (Hao et al., 2023), followed by quality control, normalization, dimensionality reduction, and clustering. Batch effects were corrected using Harmony (Korsunsky et al., 2019), and germ-cell states were annotated based on marker-gene expression. Contaminating somatic cells were removed before downstream analysis.

Chicken gene identifiers and symbols from the bulk-derived signatures were matched to genes present in the single-cell object. The induced and repressed signatures were then scored in individual cells using the Seurat AddModuleScore function. Thus, the bulk and single-cell datasets were connected through gene-set scoring rather than formal projection or deconvolution. Module scores were compared across annotated germ-cell states and visualized using violin plots.

To examine how these signatures changed along germ-cell development, pseudotime was inferred using Slingshot (Street et al., 2018), with the Harmony-corrected UMAP embedding and annotated germ-cell states used as input. PGCs were specified as the starting cluster. Module scores for the induced and repressed signatures were plotted against Slingshot pseudotime and smoothed using generalized additive models. Median pseudotime values for each annotated germ-cell state were used to indicate their relative positions along the trajectory.

### Cell-cell communication analysis

For cell–cell interaction analysis, the publicly available single-cell RNA-sequencing dataset was reprocessed as described in the previous section, and cell types were annotated following (Trost et al., 2025). Intercellular signaling was inferred from the annotated dataset using CellChat v2.2 (Jin et al., 2024). Because no chicken-specific ligand–receptor database is currently available, the human interaction database (CellChatDB.human) was used after mapping chicken gene identifiers to their one-to-one human orthologs using biomaRt. Log-normalized expression values and cell-type identities were supplied as input, and communication probabilities were computed with computeCommunProb using default parameters (trimean). Cell groups represented by fewer than 10 cells were excluded (filterCommunication, min.cells = 10). Pathway-level networks were aggregated with aggregateNet, and network centrality was computed with netAnalysis_computeCentrality. Aggregate interaction number and strength were visualized as circle plots, signaling-role contributions across pathways as heatmaps, and ligand–receptor interactions directed toward individual germline populations as bubble and chord diagrams.

### Safe-harbor candidate identification and transgene targeting

To identify candidate safe-harbor loci for transgene insertion in chicken PGCs, we analyzed non-coding RNA loci for stable and robust expression across available RNA-seq datasets. We first downloaded a large developmental transcriptomic dataset covering seven organs (brain, cerebellum, heart, kidney, liver, ovary, and testis) spanning day 10 post-conception to day 155 post-hatch, comprising 217 RNA-seq libraries (Cardoso-Moreira et al., 2019). FASTQ files were downloaded and mapped to the chicken reference genome (bGalGal1.mat.broiler.GRCg7b). Genes were ranked by mean expression, calculated on the log_10_(FPKM + 1) scale, and by coefficient of variation across samples. Candidate loci were selected from genes annotated as long non-coding RNAs that ranked within both the top 2,000 most highly expressed genes and the top 2,000 most stably expressed genes (Table S3).

Because a safe-harbor locus should ideally be transcriptionally permissive while minimizing disruption of protein-coding genes, candidate loci were further filtered based on gene structure and regulatory annotation. We prioritized loci containing at least one annotated intron and inspected chromatin-accessibility annotations from the Ensembl Regulatory Build. Based on this analysis, ENSGALG00010022677, located on chromosome 14 (Primary_assembly 14: 4,728,599–4,731,792, forward strand; bGalGal1.mat.broiler.GRCg7b, CM028495.1), was selected as a candidate safe-harbor locus (GSH). This locus is annotated as a long non-coding RNA, contains one intron, and its intronic region largely overlaps with open chromatin. The transgene cassette was targeted to this intronic region to minimize disruption of protein-coding genes while taking advantage of a transcriptionally active and accessible genomic environment.

To generate an *SYCP3*pro-EGFP reporter line, the candidate GSH on chromosome 14, corresponding to ENSGALG00010022677, was first amplified from PGC genomic DNA using primers flanking the target region (Table S6). The PCR product was verified by Sanger sequencing, and the confirmed sequence was used to design the sgRNA (Concordet & Haeussler, 2018) and the left and right homology arms. A donor vector containing the *SYCP3* promoter-driven EGFP reporter and a PGK-puromycin resistance cassette flanked by the homology arms was synthesized by VectorBuilder (donor sequence in Table S7). The sgRNA targeting the intronic region of the chromosome 14 lncRNA locus, designated sgRNA-lncR-chr14, was cloned into the pSpCas9(BB)-2A-GFP (PX458) plasmid (Addgene plasmid no. 172221; (Wu et al., 2021), and the resulting construct was confirmed by Sanger sequencing.

Female ZW PGCs were co-transfected with the sgRNA/Cas9 plasmid and the donor vector using Lipofectamine 2000 Reagent (Thermo Fisher Scientific), following the protocol described previously (Fallahshahroudi et al., 2025). At 48 h after transfection, cells were treated with an empirically determined puromycin concentration. Correct integration of the reporter cassette was assessed using genomic DNA extracted from the puromycin-resistant cells with the primers listed in Table S6 using PCR followed by Sanger sequencing.

### Induction and analysis of the SYCP3pro-EGFP reporter

The validated *SYCP3*pro-EGFP reporter PGCs were cultured under either standard growth-factor conditions or reduced growth-factor conditions containing half the standard concentrations of FGF and Activin. Cells were treated with either 100 nM RA together with 300 ng/mL BMP2, or 200 nM RA together with 600 ng/mL BMP2. Untreated cells maintained under the corresponding growth-factor conditions served as controls. EGFP fluorescence was monitored by fluorescence microscopy throughout the treatment period. At the end of treatment, cells were collected and EGFP fluorescence was quantified by flow cytometry using the same treatment groups and matched untreated controls.

### Comparative genomic investigation of ZGLP1 in avian genomes

#### Reference set and alignment

A reference set of 38 high-confidence ZGLP1/Zglp1 protein sequences spanning mammals, sauropsids (including crocodilians and the intact-ZGLP1 galliform *Lagopus muta*), amphibians, teleosts and cartilaginous fish was compiled from NCBI RefSeq (accessions in Table S5). Sequences were aligned with MAFFT L-INS-i (--localpair --maxiterate 1000) to produce a high-accuracy multiple sequence alignment for HMM construction and for the Fig. 5B panel.

#### Profile HMM construction

Three profile hidden Markov models were built from the multiple sequence alignment with HMMER v3.4 hmmbuild, spanning successively more conserved regions: (i) a full-length model (333 aligned positions), (ii) a mid-column model (281 positions after trimming columns with gap fraction ≥ 0.3), and (iii) a core zinc-finger model (80 positions spanning the canonical CASCxT…CGIR…CxxCW GATA-type zinc-finger). All three HMMs were used in parallel to maximise sensitivity to divergent or truncated relicts.

#### Genome and proteome searches

HMM searches (hmmsearch, phmmer) and *Lagopus*-anchored tblastn searches (BLAST+ 2.16.0; queries: *Lagopus muta* XP_048788870.1 and XP_039910342.1) were performed against three chicken assemblies — bGalGal1.mat.broiler.GRCg7b, Red Jungle fowl GRCg6a (GCF_000002315.6) and telomere-to-telomere Huxu bGalGal2b (GCA_024206055.2) — and against seven additional avian assemblies: *Lagopus muta* (GCF_023343835), *Meleagris gallopavo* (GCF_905368555, GCF_047663525), *Coturnix japonica* 2025 WE-NBRP24 (GCF_054131305.1) and 2016 WashU (GCA_001577835.2), *Pavo cristatus* (GCF_048771995), *Anas platyrhynchos* (GCF_036370855), *Dromaius novaehollandiae* (GCF_030867095) and *Taeniopygia guttata* (GCF_003957565.2). Candidate loci were validated by synteny with seven conserved flanking anchors (ICAM5-like, ICAM5, FDX2, RAVER1, TYK2, CDC37, PDE4A) and, where tblastn returned only partial hits, by six-frame conceptual translation of the genomic region using EMBOSS transeq.

#### Diagnosis of pseudogenization

Chicken chr30 loci were classified as pseudogenized when they satisfied all of (i) recovery at the canonical syntenic position, (ii) presence of in-frame stop codons in the *ZGLP1* reading frame, (iii) low-complexity expansion inflating the locus relative to the intact Lagopus muta ortholog (chicken chr30 locus ∼10 kb vs ∼3.2 kb in Lagopus; best BLASTN identity 72%), (iv) proline enrichment over the downstream 200-aa window (∼33% proline), and (v) N-content close to zero (0% N in T2T Huxu, 0.2% N in GRCg6a), confirming that the detected stops are real sequence rather than assembly gaps. Best-hit tblastn coordinates and statistics for all species surveyed are compiled in Table S5: T2T Huxu CP100584.2:149,802–152,582 (E = 1.70×10⁻⁴³, 53.4% identity over 251 aa); GRCg6a NC_028739.2:219,654–222,480 (E = 4.89×10⁻⁴⁰, 53.5% over 273 aa); *Lagopus muta* NW_026040201.1:28,163–30,712 (E = 2.64×10⁻¹³⁸, 89.4%); *Meleagris gallopavo* NW_028147023.1:15,513–17,274 (E = 3.46×10⁻⁸⁸, 76.6% over 295 aa); *Coturnix japonica* GCA_001577835.2 KQ967012.1 (zinc-finger halves at E = 6.22×10⁻⁹, 96.6% identity and E = 2.07×10⁻⁵, 68.3%); *Taeniopygia guttata* NC_133062.1:985,690–985,887 (E = 1.48×10⁻⁶, 54.8% over 93 aa).

#### Coturnix cross-verification

The current Coturnix japonica RefSeq assembly (GCF_054131305.1, WE-NBRP24, 2025; long-read) initially indicated two in-frame stops in the second half of the *ZGLP1* zinc finger. This region was cross-verified against the independent 2016 short-read WashU assembly (GCA_001577835.2) by tblastn of *Lagopus muta* XP_048788870.1 into both genomes. The 21/120 bp of DNA-level differences between the two assemblies in the affected 40-aa region were concentrated in a poly-C/poly-G stretch — the characteristic homopolymer-indel error signature of long-read sequencing — and the WashU 2016 sequence, which recovers both halves of the ZGLP1 zinc finger in a single open reading frame with no in-frame stops, was used for the Fig. 5B alignment. A reconstructed *Coturnix japonica* ZGLP1 zinc-finger sequence (SCASCKTQQTPLWRAAENGTPLCNACGIRYRKYRRRCRRCWSVPRRSRAPPPLRCPQCGDEAPHVGGGINMDPPRS; 76 aa; concatenation of the two zinc-finger halves with the single-residue MSA gap collapsed) is provided in Supplementary Data 1.

## Data Availability

The raw sequencing data generated in this study have been deposited in the European Nucleotide Archive (ENA) under project accession PRJEB111572. All ZGLP1 sequence data analyzed in this study were obtained from publicly available RefSeq and GenBank assemblies, with accession numbers provided in Table S5 and File S1. File S1 contains the FASTA sequences used to generate the multiple sequence alignment shown in Fig. 5B. Table S5 (Supplementary_Table_5_ZGLP1.xlsx) contains the ZGLP1 detection status for each species in Sheet S1 and the zinc-finger sequences and associated metadata in Sheet S2.

## Acknowledgments

This study was supported by grants from the Swedish Research Council for Sustainable Development (Formas; 2023-01396) and the Carl Trygger Foundation (CTS 24:3578) to S.Y.T., and by grants from the Swedish Research Council (VR; 2025-05129), the Swedish Research Council for Sustainable Development (Formas; 2021-00513), and the Carl Trygger Foundation (CTS 24:3571) to A.F. Claude Opus 4.8 (Anthropic) and ChatGPT GPT-5.5 (OpenAI) were used for language editing and refinement of the manuscript text, and for optimisation and debugging of custom analysis code. All scientific content, data interpretation, and conclusions are the sole responsibility of the authors.

## Author contributions

M.H.G. performed experiments and computational analyses and contributed to manuscript writing. S.C. and A.N. generated the SYCP3pro-EGFP reporter line. C.G., S.D., and Y.X. contributed to the establishment and optimization of chicken PGC culture. M.M.A. supported microscopy and imaging experiments. C.M. performed the evolutionary analysis. M.J. performed the bioinformatic analysis used to identify the candidate genomic safe-harbor locus. A.F. and S.Y.T. conceived and supervised the study, secured funding, and wrote the manuscript with input from all authors. All authors reviewed and approved the final manuscript.

## Supplementary materials

**Table S1. Differential expression of genes in RA-treated versus control chicken PGCs.** Columns give the Ensemble gene ID (*ensembl_gene_id*); mean normalized expression (*baseMean*); the log_2_ fold change and its standard error (*log_2_FoldChange*, *lfcSE*); the Wald statistic (*stat*); and the raw and Benjamini– Hochberg-adjusted *P* values (*pvalue*, *padj*; genes called differentially expressed at *padj* < 0.05); gene biotype and symbol (*external_gene_name*). The final two columns (*ovary_log_2_FoldChange*, *ovary_padj*) give the corresponding values from the chicken ovary developmental analysis (Cardoso-Moreira et al., 2019) for direction-matched comparison.

**Table S2. Differential expression of genes in RA and BMP2-treated versus control chicken PGCs.** Columns give the Ensemble gene ID (*ensembl_gene_id*); mean normalized expression (*baseMean*); the log_2_ fold change and its standard error (*log_2_FoldChange*, *lfcSE*); the Wald statistic (*stat*); and the raw and Benjamini–Hochberg-adjusted *P* values (*pvalue*, *padj*; genes called differentially expressed at *padj* < 0.05) gene biotype and symbol (*external_gene_name*). The final two columns (*ovary_log_2_FoldChange*, *ovary_padj*) give the corresponding values from the chicken ovary developmental analysis for direction-matched comparison.

**Table S3. Stably expressed lncRNAs in the chicken PGC transcriptome.** Long non-coding RNAs (Ensembl *Gallus gallus* annotation) ranked by expression stability across *n* = 9 samples. Columns: ensembl_gene_id, gene accession; seqnames, chromosome/scaffold (JAENSK0* = unplaced contig); start**/**end**/**width, genomic coordinates and length (bp); strand, orientation; gene_name, symbol where assigned (NA = none); gene_biotype, Ensembl biotype; mean_expr**/**sd_expr, mean and SD of normalized expression [unit]; cv_expr, coefficient of variation (stability metric, lower = more stable); mean_rank, rank by mean expression; cv_rank, rank by CV; n_exons, exon count. Ordered by ascending cv_expr.

**Table S4. Inferred ligand–receptor interactions across ovarian populations.** All 3,373 significant interactions inferred with CellChat (Human ligand–receptor database; <10 cells per population excluded), spanning 10 cell populations and 71 pathways. Columns give source and target population, ligand, receptor, communication probability (*prob*), permutation *P* value (*pval*), interaction and pathway names, interaction category (*annotation*), and database reference (*evidence*).

**Table S5. In silico search for a chicken ZGLP1 orthologue and its status in additional galliform sister taxa**. The table lists per-species assembly identifiers, best-hit contigs and coordinates, tblastn E-values, percent identities, and inferred status (intact, relict-pseudogene, assembly-truncated, or germline-restricted) for ten species surveyed by HMM- and Lagopus-anchored tblastn. Three chicken assemblies (bGalGal1.mat.broiler.GRCg7b [assembly-truncated at chr30 position 755,666 bp], GRCg6a and T2T Huxu bGalGal2b) recover a pseudogenized ZGLP1 relict on chromosome 30, while Lagopus muta, Meleagris gallopavo and Coturnix japonica (2016 WashU assembly, cross-verified against the 2025 WE- NBRP24 assembly) retain an intact ZGLP1 coding sequence.

**Table S6. Oligonucleotides used for sequencing, sgRNA cloning, and genomic validation of the chromosome 14 lncRNA locus.** Oligonucleotide sequences are shown in the 5′–3′ direction. Lowercase nucleotides in the sgRNA oligonucleotides indicate BbsI-compatible overhangs used for cloning. Expected PCR product sizes are provided for the corresponding primer pairs. F, forward; R, reverse; sgRNA, single-guide RNA; bp, base pairs.

**Table S7. DNA constructs used for CRISPR–Cas9-mediated homology-directed repair in chicken PGCs.** The pSA-1000 donor construct contained the *SYCP3* promoter–EGFP reporter and PGK promoter– puromycin-resistance cassette flanked by left and right homology arms targeting the chromosome 14 lncRNA locus. pX458_lncR-Chr14 was generated by cloning the Chr14 lncRNA-targeting sgRNA into 2X_pX458_pSpCas9(BB)-2A-GFP (Addgene plasmid #172221; Wu et al., 2021). hU6, human U6 promoter; sgRNA, single-guide RNA; Cas9, CRISPR-associated protein 9; EGFP, enhanced green fluorescent protein; PGK, phosphoglycerate kinase promoter; CAG, CMV early enhancer/chicken β-actin promoter; T2A, *Thosea asigna* virus 2A self-cleaving peptide; PuroR, puromycin-resistance gene; bGH polyA, bovine growth hormone polyadenylation signal; SV40 polyA, simian virus 40 polyadenylation signal; LHA, left homology arm; RHA, right homology arm; HDR, homology-directed repair.

**File S1. ZGLP1 zinc-finger sequences used for the multiple sequence alignment.** FASTA of 33 unaligned ZGLP1 zinc-finger amino acid sequences (76–105 aa) used as input for the Fig. 5B MSA, comprising 25 orthologues from the HMM reference panel (by NCBI accession) plus eight descriptive-label sequences — the two chicken chr30 relict conceptual translations, the three intact-ZGLP1 galliform sisters (Lagopus, Meleagris, reconstructed Coturnix), the Dromaius ratite outgroup, and a duplicated Lagopus anchor query.

